# Enhancement of antibody thermostability and affinity by computational design in the absence of antigen

**DOI:** 10.1101/2023.12.19.572421

**Authors:** Mark Hutchinson, Jeffrey A. Ruffolo, Nantaporn Haskins, Michael Iannotti, Giuliana Vozza, Tony Pham, Nurjahan Mehzabeen, Harini Shandilya, Keith Rickert, Rebecca Croasdale-Wood, Melissa Damschroder, Ying Fu, Andrew Dippel, Jeffrey J. Gray, Gilad Kaplan

## Abstract

Over the last two decades, therapeutic antibodies have emerged as a rapidly expanding domain within the field biologics. In silico tools that can streamline the process of antibody discovery and optimization are critical to support a pipeline that is growing more numerous and complex every year. In this study, DeepAb, a deep learning model for predicting antibody Fv structure directly from sequence, was used to design 200 potentially stabilized variants of an anti-hen egg lysozyme (HEL) antibody. We sought to determine whether DeepAb can enhance the stability of these antibody variants without relying on or predicting the antibody-antigen interface, and whether this stabilization could increase antibody affinity without impacting their developability profile. The 200 variants were produced through a robust highthroughput method and tested for thermal and colloidal stability (T_onset_, T_m_, T_agg_), affinity (K_D_) relative to the parental antibody, and for developability parameters (non-specific binding, aggregation propensity, self-association). In the designed clones, 91% and 94% exhibited increased thermal and colloidal stability and affinity, respectively. Of these, 10% showed a significantly increased affinity for HEL (5-to 21-fold increase), with most clones retaining the favorable developability profile of the parental antibody. These data open the possibility of *in silico* antibody stabilization and affinity maturation without the need to predict the antibody-antigen interface, which is notoriously difficult in the absence of crystal structures.

## Introduction

Antibodies are well-suited for a wide range of therapeutic and diagnostic applications due to their ability to bind targets with high affinity and specificity. However, despite their utility, the development of antibodies against new targets remains an expensive and time-consuming endeavor. Traditionally, antibodies have been discovered using hybridoma technology [1] or selected from large sequence libraries [2]. Although these approaches are effective at uncovering target-specific antibodies, the antibodies they produce often require extensive optimization before they are fit for practical use [3, 4]. Among the properties that are typically subject to optimization are affinity, thermal stability, aggregation propensity, and solubility. Optimization of these attributes can be achieved through rational protein engineering or library screening approaches, but this potentially requires many rounds of trial and error to obtain a well-behaved molecule. Computational methods for antibody optimization promise to accelerate this process, through suggestion of mutations and prediction of mutational effects on key attributes.

The biophysical properties of antibodies are determined in large part by their molecular structure. As such, computational assessment of these properties requires an accurate threedimensional structure. While experimental structure determination has long been a low-throughput, expensive undertaking, computational prediction of protein structures has significantly improved in accuracy in recent years. Most notably, the AlphaFold2 system has demonstrated high-quality structure prediction across a broad range of protein families [5], rivaling experimental accuracy. In parallel with the development of AlphaFold2 and successor methods [6-8], a set of antibody-specific structure predictors has demonstrated state-of-the-art accuracy with significantly less computational overhead [9-11]. One such method, DeepAb, approaches antibody structure prediction through learning a set of inter-residue geometric potentials [11]. These potentials compose a learned pairwise energy function that is subsequently minimized through folding simulations to produce a three-dimensional structure of the antibody. Subsequent methods have eschewed with this latter step, opting instead to predict the antibody structure directly in an end-to-end fashion [9, 10, 12].

As machine learning methods have become increasingly accurate at mapping between sequence and structure, they have also enabled new methods for protein design. Referred to as “hallucination,” these methods aim to explore sequence space in search of an input that optimizes the predicted structure [13]. Early hallucination methods utilized Markov chain Monte Carlo sampling techniques to accept or reject mutations according to a structure predictors confidence [13]. Subsequent methods have utilized structure predictors as differentiable protein designers [14], directly optimizing the input sequence according to an objective function – e.g., adopting a particular fold, hosting a scaffold, or binding another protein. In the antibody design space, similar methods have utilized DeepAb for designing libraries of putative binders or optimizing the framework residues [15]. Beyond repurposing of structure predictors, several other modeling strategies have shown promise for antibody design. Protein language models are a class of self-supervised machine learning models that learn directly from large databases of protein sequences [16, 17]. Numerous antibody-specific language models have been trained on immune repertoire data (encompassing ~0.5 billion antibody sequences), enabling them to learn the underlying rules of antibody sequences without need for structural or functional information [18, 19]. Applications of these models include structure prediction [10], humanization [20], and library generation [17, 21]. Score-based (or diffusion) models are another class of models that have shown promise for antibody design. These models are trained to iteratively generate antibody structures, typically for redesigning or optimizing the binding interface with the antigen [22]. However, despite the considerable interest in antibody design methods, many methods remain unvalidated in the experimental setting. Going forward, it will be particularly important to assess the effectiveness of tools to best inform future directions.

The amount of data needed to train and validate certain computational techniques is extremely large. A recent publication described the development of ProGen, a language model that can design protein sequences with predicted biological function, using 280 million protein sequences from existing databases [23]. Likewise, the thermostability predicting TemStaPro was trained using >2 million protein sequences from organisms with known optimal growth temperatures from existing databases [24]. Synthetic training exercises suggest that at least 10^4^ antibody-antigen affinity measurements will be needed to train a reliable machine learning model [25]. Current semi or fully automated highthroughput (HT) experimental methods cannot produce and test such large numbers of biologics but may produce 10^2^-10^3^ clones per production cycle, allowing for the generation of large datasets over time. To support HT biologics production, methods for HT cloning, transfection and purification need to be implemented [26-28]. In addition, HT experimental methods are required to determine key attributes of the produced biologics, such as expression titers, monomer content, and antigen specificity [29-31]. Such methods must require small amounts of material and utilize 96 or 384-well plates for sampling.

To complement and support these HT experimental techniques, bioinformatic tools supporting each of these stages are necessary. HT cloning requires tools that allow in silico molecule design, automated vector map creation and automated deconvolution of sequencing files for the designed molecules. The HT protein expression, purification, and characterization requires automated liquid handling and colony picking systems and full tracking of all associated data in a laboratory information management system (LIMS) or biologics database [30]. The combination of automated liquid handling systems, HT experimental techniques and enabling bioinformatics tools allow for a fully automated HT biologics workflow. Such an end-to-end automated workflow for the cloning, production and characterization of multi-specifics has been described in detail [30]. Here we describe a semi-automated workflow using manual HT cloning and a semi-automated transfection/purification workflow that we used to produce the computationally designed variants in this study as a step towards full process automation.

In this work, we present a strategy for optimization of antibody thermostability and affinity using DeepAb [11] as well as the experimental framework to validate the optimized variants. Starting from a natural antibody against hen egg lysozyme and guided by existing deep mutational scanning affinity data, we designed variants by optimizing the structure prediction confidence. Experimental validation demonstrates that this design strategy is highly effective at improving stability and affinity.

## Results

### Computational design of optimized anti-HEL antibody variants

To design optimized antibody variants, we adopted the mutation ranking protocol proposed below alongside the DeepAb antibody structure prediction model [11]. DeepAb is a machine learning model for antibody Fv structure prediction that formulates structure modeling as a prediction of a set of geometric potentials between pairs of residues. To produce an antibody structure these potentials are treated as an energy function and minimized in Rosetta. Additionally, Ruffolo et al. proposed to use the sharpness (or confidence) of these potentials as a proxy for mutational fitness. Computational assessment of this strategy showed promise for identifying optimized variants of three antibodies against diverse targets (hen egg lysozyme (HEL), VEGF, and QSOX1) [11]. We extend this methodology to design and experimentally test new variants of an anti-HEL antibody, as illustrated in Figure 1A.

**Figure 1.**
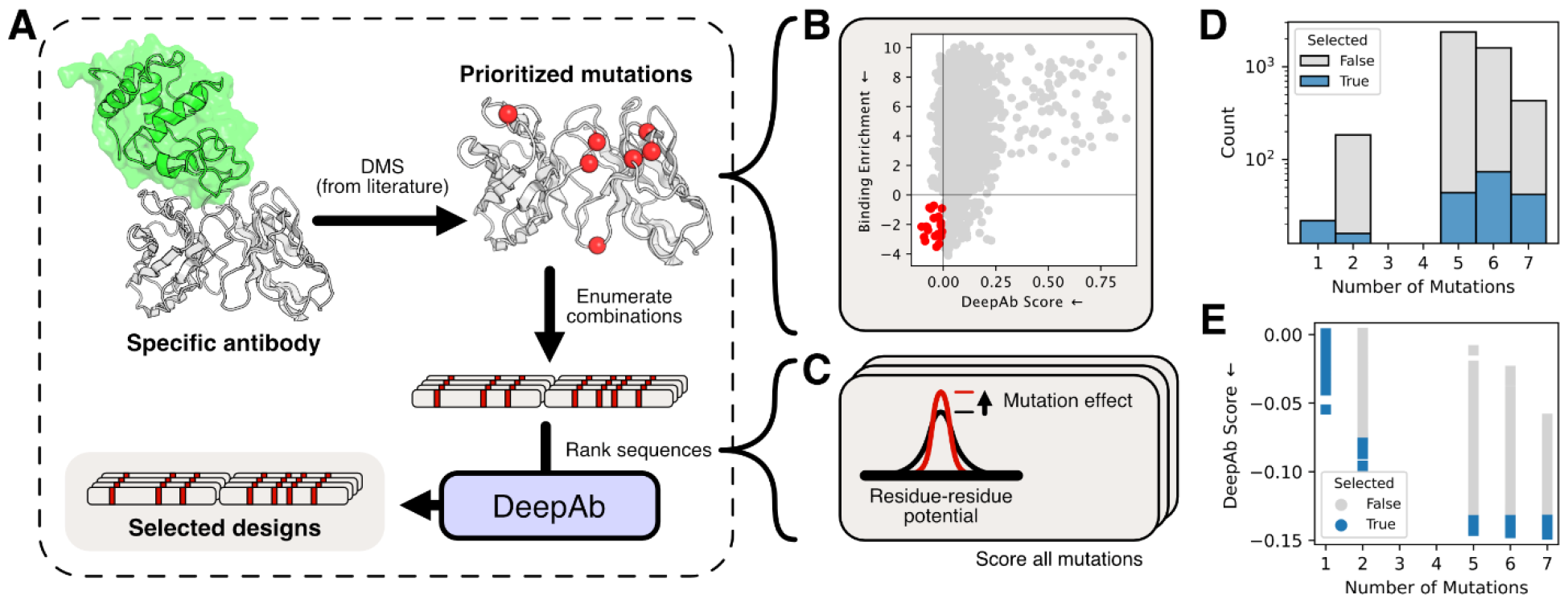
Computational design of optimized anti-HEL variants. (A) Diagram of design process using DeepAb score to identify optimal variant sequences. (B) Relationship between DeepAb score and binding enrichment for deep mutational scan data from Warszawski et al. [32]. Red points are predicted to improve over parental anti-HEL and have improved binding enrichment. (C) Illustration of increased confidence of DeepAb inter-residue potentials. Sharper potentials indicate a favorable mutation (lower score). (D) Distribution of selected variants for each number of mutations. (E) DeepAb score of selected and non-selected variants for each mutational bin.

Our design process began with deep mutational scanning (DMS) data collected by Warszawski et al [32] for an anti-lysozyme antibody (clone D44.1). The DMS data encompasses saturating mutations at 135 positions including the CDR loops, the heavy- and light-chain interface, and peripheral positions. Following Ruffolo et al., we scored these mutations relative to the parent antibody using the ΔCCE metric from DeepAb [11], which measures a change in structure prediction confidence upon mutation (Figure 1C). We refer to this metric as the DeepAb score and plot it against the experimental binding enrichment values for variants of the same anti-lysozyme antibody from Warszawski et al. [32] (Figure 1B, lower is better for both metrics). Twenty-two mutations at seven positions were predicted to improve fitness and saw an improvement in binding enrichment in Warszawski et al.’s [32] prior DMS screens (Figure 1B, red points and Figure 2). We focused on recombining these mutations to design optimized antibodies.

**Figure 2.**
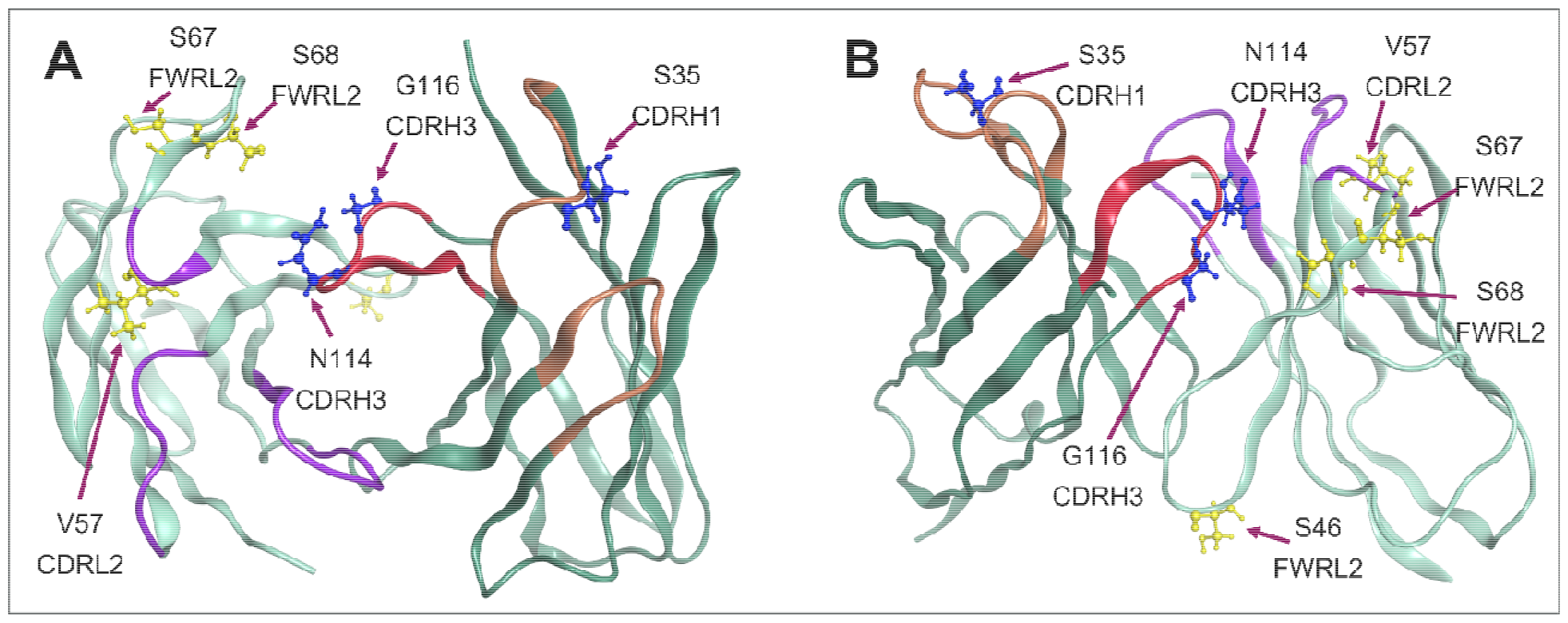
Structure of parental α-HEL Fab D44.1 with mutated positions designed using DeepAb marked. (A) “Top view” and (B) “side-view” of the D44.1 Fv marked with the positions predicted to improve antibody properties in the designed variants. Designed variants contain single, double, or combinations of 5-7 amino acid changes in these positions. CDRs H1 and H2 (orange), CDRH3 (red) and CDRs L1, L2 and L3 (purple) are marked. CDRH3 residues N114 and G116 are found in the VH:VL interface. Structure taken from PDB ID 1MLC [33]. In the VH, mutations occur in the CDRs at S35 (CDRH1), N114 (CDRH3) and at G116 (CDRH3). In the VL, mutations occur in the frameworks at positions S46 (FWRL2), S67 (FWRL3) and S68 (FWRL3), and in the CDRs at position V57 (CDRL2). Numbering and CDR annotations correspond to IMGT conventions.

We designed a large pool of anti-HEL variants by enumerating double mutations and combinations of five or more mutations. Including the original single-point mutations, our initial pool included 4,602 antibody sequences. From this pool, we selected all the single-point variants (22 total), 16 double mutants, and 160 sequences with five or more mutations (Figure 1D) according to DeepAb score (Figure 1E). We additionally included the parent anti-HEL antibody and the top design from Warszawski et al. [32] (D44.1_des_), for a total of 200 sequences (designated “Top 200”) to advance to experimental characterization (Table S1 and Figure S1).

### Manual high-throughput cloning yields correctly assembled IgG clones

Manual high-throughput (HT) cloning was achieved by conducting the cloning, transformation, plating, and sequencing steps in a 96-well plate format (Figure 3). This allowed us to obtain the desired 200 clones rapidly and with much less effort than standard cloning techniques, which use single tubes and plating each transformed clone onto a separate agar plate. All mAbs were cloned as human IgG1 in a bicistronic expression vector that encodes both the light and heavy chains of the antibody. To achieve HT cloning, synthetic DNA fragments encoding the VH and VL regions were ordered with the required complementary Gibson cloning overhangs in 96-well plates (Figure 3). The variable region fragments were then added to a plate containing a mix of the digested vector backbone, a “middle fragment” generated by PCR, and Gibson reaction mix to perform a four-piece Gibson assembly. Here, the “middle fragment” encodes the light chain constant region, polyA signal, and the promoter and signal peptide for the heavy chain expression cassette. The resulting Gibson reaction was transformed into chemically competent E. coli formatted in 96-well plates. The transformed cultures were then stamped onto large rectangular agar plates to produce single, pickable colonies. The resulting colonies were hand-picked and grown in 96-well deep well culture plates for subsequent 96-well minipreps, Sanger sequencing, and production of glycerol stocks plates. The continual use of reagents in 96-well format significantly reduces the time and effort required to clone hundreds of clones compared to standard cloning and plating techniques.

**Figure 3.**
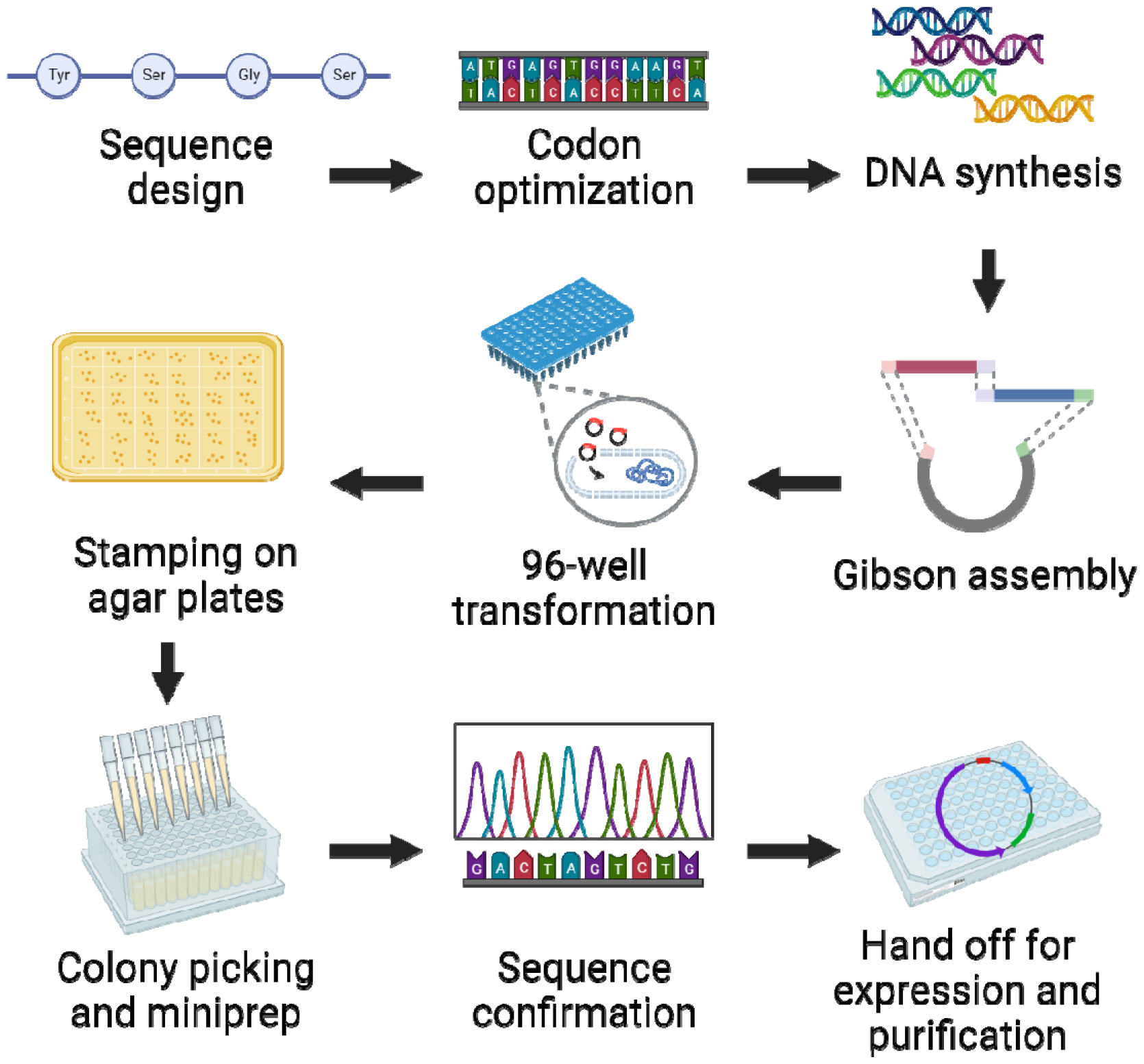
High throughput (HT) cloning workflow. Figure illustrating the steps allowing the HT cloning of several hundred clones in one experimental cycle without the use of automation. All steps (DNA synthesis, Gibson cloning, transformation, colony growth, DNA mini-prep and sequencing) were conducted in a single step manner in 96-well plate format. Figure created with BioRender.

While the first round of HT cloning resulted in only ~50% correctly assembled products, the process was improved over several iterations to optimize the fragment assembly, reduce mutagenesis rate, and increase plasmid yield. To overcome the low initial success rate, several modifications were made to the process. First, different competent cells were tested for transformation efficiency and plasmid yield. While NEB 10-beta and TOP10 cells showed similar transformation efficiency, TOP10 cells produced higher plasmid yields and were chosen for compatibility with downstream protein production. Next, to reduce the mutagenesis rate in plasmid assembly, we changed from using Gibson Assembly Master Mix (NEB) to NEBuilder HiFi DNA Assembly Master Mix. We found that this reaction mix produced far fewer mutations around the sites of ligation, which allowed us to sequence fewer colonies for correct assembly. In general, we found that mutations were introduced almost exclusively in the VH and VL due to the inherent error rate of gene fragment synthesis (reported median error rate of 1:5000 bp). Finally, we optimized the amount of post-reaction assembled Gibson mixture used in transformation. We found that diluting the assembly reaction 1:1 with water prior to transformation ultimately led to more colonies. These changes cumulatively resulted in a cloning success rate of >90% per cycle. Utilizing these new techniques allowed us to go from receiving gene fragments to having sequence confirmed plasmids in 4 days. Compared to standard methods this HT process reduces hands-on time significantly and generates much less waste (single agar plates vs single stamped plate, TOP10-cell tubes vs plates, and overall picking of fewer colonies to obtain the correct sequence). Additionally, this HT process is highly amenable to automation with the addition of liquid handling and colony picking robotic systems at various stages.

### HT semi-automated expression and purification of computationally designed antibody variants produces high yields of purified protein

Expression and purification of the parental D44.1 IgG and 199 designed variants were conducted using a semi-automated expression and purification scheme (Figure 4 A-F). Briefly, Expi293 cells were transiently transfected with sequence-confirmed expression plasmids, grown in 24-well deep well plates and purified from the supernatants using magnetic protein A beads. Figure 5-A shows the yields achieved across four different expression batches. To achieve the improvement in yields seen in Figure 5-A, two expression parameters were optimized. Between batches one and two, the schedule and amounts of proprietary cell culture feed used were optimized. Between batches two and three the quality of the transfecting DNA was improved by a combination of changing the DNA purification method from a bead-based method to a column-based purification method and stringent DNA QC. DNA QC focused on getting high quality super-coiled DNA of the correct size without any observable “laddering” upon visualization. These changes led to an increase in the mean yield from 0.16 mg (+/-0.17) (batch A) to 1.13 mg (+/-0.24) (batch D) (Figure 5-A). After purification, the IgGs underwent QC by

**Figure 4.**
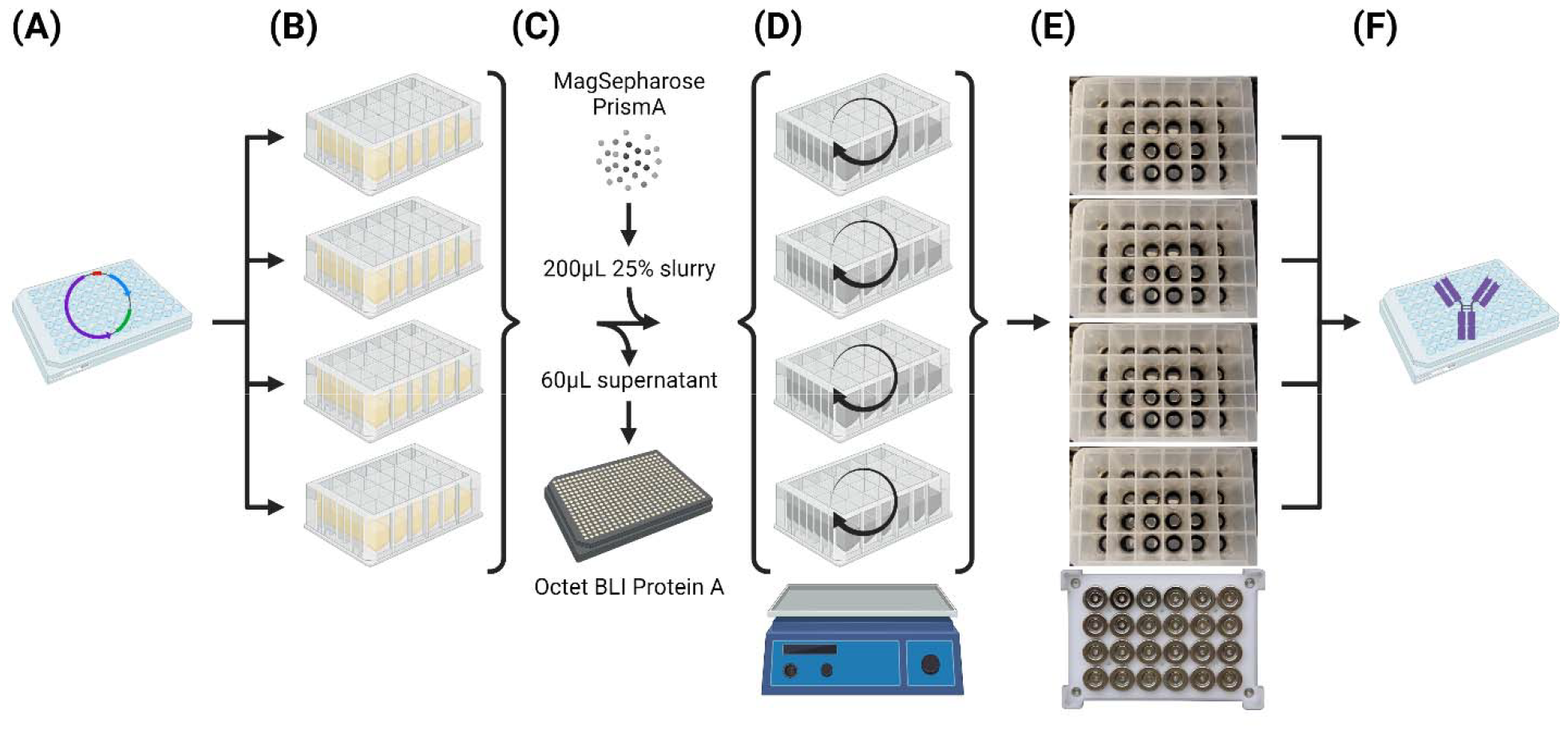
High-throughput antibody expression and purification. (A) In 96-well format, DNA was normalized to match concentrations prior to transfection. (B) Expi293 cells (3 mL) were dispensed in 24-deep-well plates and transfected with DNA from the 96-well plate in A1, A2, B1, B2 quadrant format. (C) Seven days after transfection, cells were centrifuged and supernatant transferred to new 24-deep-well plates. An aliquot of supernatant (60 µL) was removed to measure antibody titer by Octet BLI Protein A prior to the addition of 200 µL 25% slurry (i.e., 50 µL settled) of MagSepharose PrismA Protein A beads. (D) Supernatant and Protein A beads were incubated with shaking at room temperature for 4 hours to immobilize antibodies on beads. (E) Using a 24-well plate magnet, supernatant was removed, Protein A beads with bound antibodies were washed, and antibodies were eluted. (F) Antibodies were then transferred back into 96-well format to the appropriate A1, A2, B1, B2 quadrants from which their DNA originated. Figure created with BioRender.

**Figure 5.**
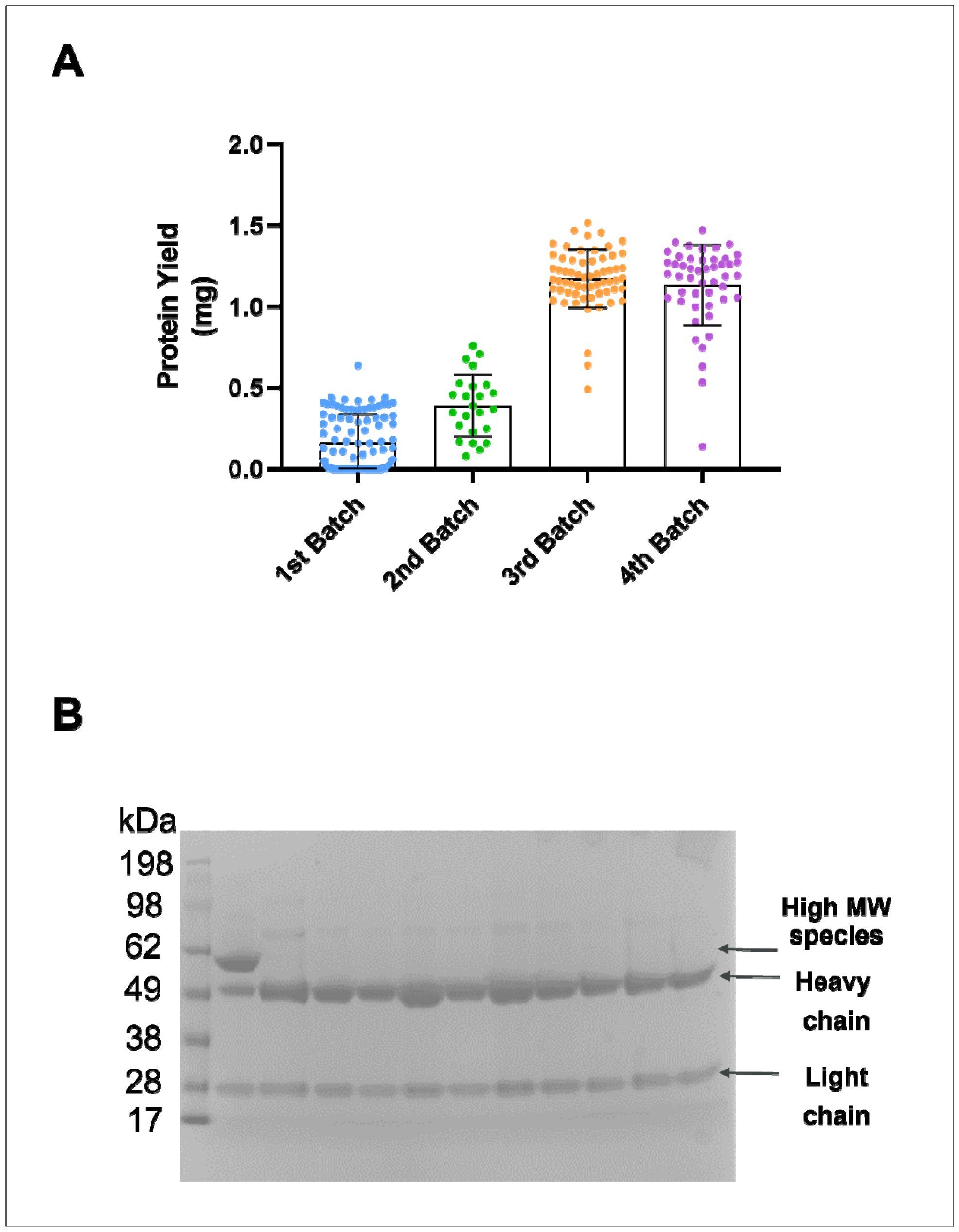
D44.1 variant antibody yields and SDS-PAGE analysis. (A) Purified IgG Yields from the 4 separate protein production cycles. Yields were 0.16 mg (+/-0.17), 0.39 mg (+/-0.19), 1.17 (+/-0.17) and 1.13 mg (+/-0.24) for purification batches 1, 2, 3, and 4 respectively. (B) Reduced SDS-PAGE of representative purified IgGs shows correct heavy and light chain banding of IgG with minimal impurities. Approximately 5% of the produced mAbs showed a high molecular weight species and needed to be reexpressed and purified.

SDS-PAGE (Figure 5-B). By SDS-PAGE analysis only 5% of clones (10/200) showed high molecular weight (HMW) species and were re-expressed and re-purified.

### Binding kinetics of parental D44.1_IgG_ and D44.1_Fab_ to HEL

As part of the validation of the experimental setup before screening the designed variants for binding to HEL, a preliminary binding experiment was run between parental D44.1 and HEL using biolayer-interferometry (BLI) (Figure 6 A-D). In this experiment, parental D44.1_IgG_ or D44.1_Fab_ was loaded onto anti-human FAB2G Octet probes and associated with varying concentrations of HEL (Figure 6 A-B). The measured IgG K_D_ and Fab K_D_ were 6 nM and 3.75 nM respectively (Figure 6 A-B), which is a significantly higher affinity than some published D44.1 affinity values of around 137 nM [32, 34], but in line with other publications [35-37] (measurement techniques vary between publications). To verify that the observed difference in our measurements vs. some of the literature reported values were not method related, we repeated the measurement using a similar experimental setup but this time using surface plasmon resonance (SPR) (Figure 6 E-F). The measured affinity (K_D_) for D44.1_IgG_ and D44.1_Fab_ was 0.93 nM and 1.19 nM respectively (Figure 6 E-F), in agreement with our BLI measurements (Figure 6 A-B), indicating that using SPR or BLI is not the source of the difference from some of the previously reported affinities for D44.1.

**Figure 6.**
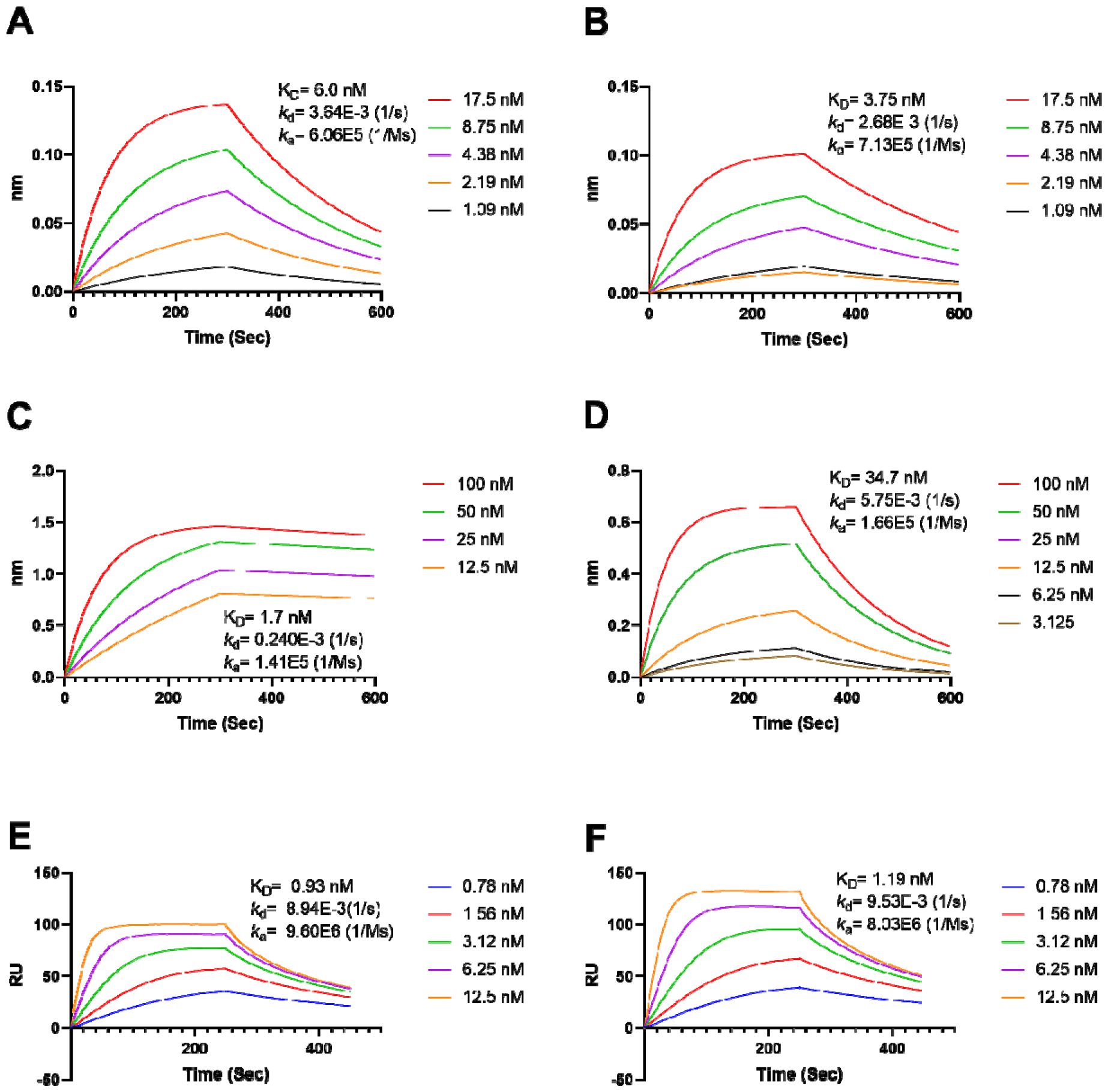
Binding kinetics of parental D44.1_IgG_ and D44.1_Fab_ to HEL. Binding kinetics for D44.1_IgG_ and D44.1_Fab_ to HEL were measured by BLI (A-D) or SPR (E-F). **A-B** BLI kinetics measurement using D44.1_IgG_ (A) or D44.1_Fab_ (B) loaded onto anti-CH1 Octet probes and dipped into varying concentrations of HEL. Measured affinity (K_D_) was 6 and 3.75 nM for D44.1_IgG_ and D44.1_Fab_ respectively. **C-D** BLI kinetics measurement for Biotinylated-HEL loaded onto anti-streptavidin Octet probes dipped into varying concentrations of D44.1_IgG_ (C) or D44.1_Fab_ (D). Measured affinity (K_D_) using this experimental setup was 1.7 and 34.7 nM for D44.1_IgG_ and D44.1_Fab_ respectively. **E-F** SPR kinetics measurement using D44.1_IgG_ (E) or D44.1_Fab_ (F) loaded CM5 chips and associated with varying concentrations of HEL. Measured affinity (K_D_) using this experimental setup was 0.93 and 1.19 nM for D44.1_IgG_ and D44.1_Fab_ respectively.

To test if our binding results were being affected by avidity effects, streptavidin coated Octet probes were loaded with biotinylated HEL and associated with either D44.1_IgG_ or D44.1_Fab_ (Figure 6 C-D). In this experimental setup, a similar affinity (IgG K_D_) of 1.7 nM was observed for D44.1_IgG_, which is unsurprising as this experimental orientation with bivalent IgG would lead to avidity binding effects. In contrast, D44.1_Fab_ exhibited an affinity (Fab K_D_) of 34.7 nM (Figure 6 D), indicating that we were possibly seeing avidity effects in our measurements using probe/chip loaded D44.1_IgG_ or D44.1_Fab_ (Figure 6 A-B, E-F). This observed effect may be caused by using antigen that is not fully monomeric in the binding experiments. To verify the monomer content of the commercial HEL we were using, we analyzed commercial HEL from two different sources by UPLC-SEC. As shown in Figure S2, both commercial batches showed main peaks of the correct approximate size, indicating monomeric antigen. It is therefore unclear what is causing the differences in the literature reported affinities for D44.1 and the difference in measured affinity between in-solution D44.1_Fab_ (Figure 6 D) and all other experimental orientations (Figure 6 A-B;E-F).

Based on these results, we decided that primary screening for affinity improvements would proceed with a BLI based binding screen using the produced 200 variants in IgG format (IgG K_D_). A select panel of binders would then be fragmented into Fabs and tested for in-solution affinity (Fab K_D_) at a second stage.

### “Top 200” designed variants exhibit greater thermostability and affinity

The thermostability and colloidal characteristics of the 200 purified D44.1 variants in IgG format (designated “Top 200”) were analyzed using differential scanning fluorimetry (DSF) and static light scattering (SLS). We focused here on the T_m1_ measurement as a proxy for overall thermostability, alongside the aggregation temperature (T_agg_) and unfolding onset temperature (T_onset_) (Figure 7, Table S1). The IgG variants were then screened for improved affinity by BLI (IgG K_D_) (Figure 7, Table S1).

**Figure 7.**
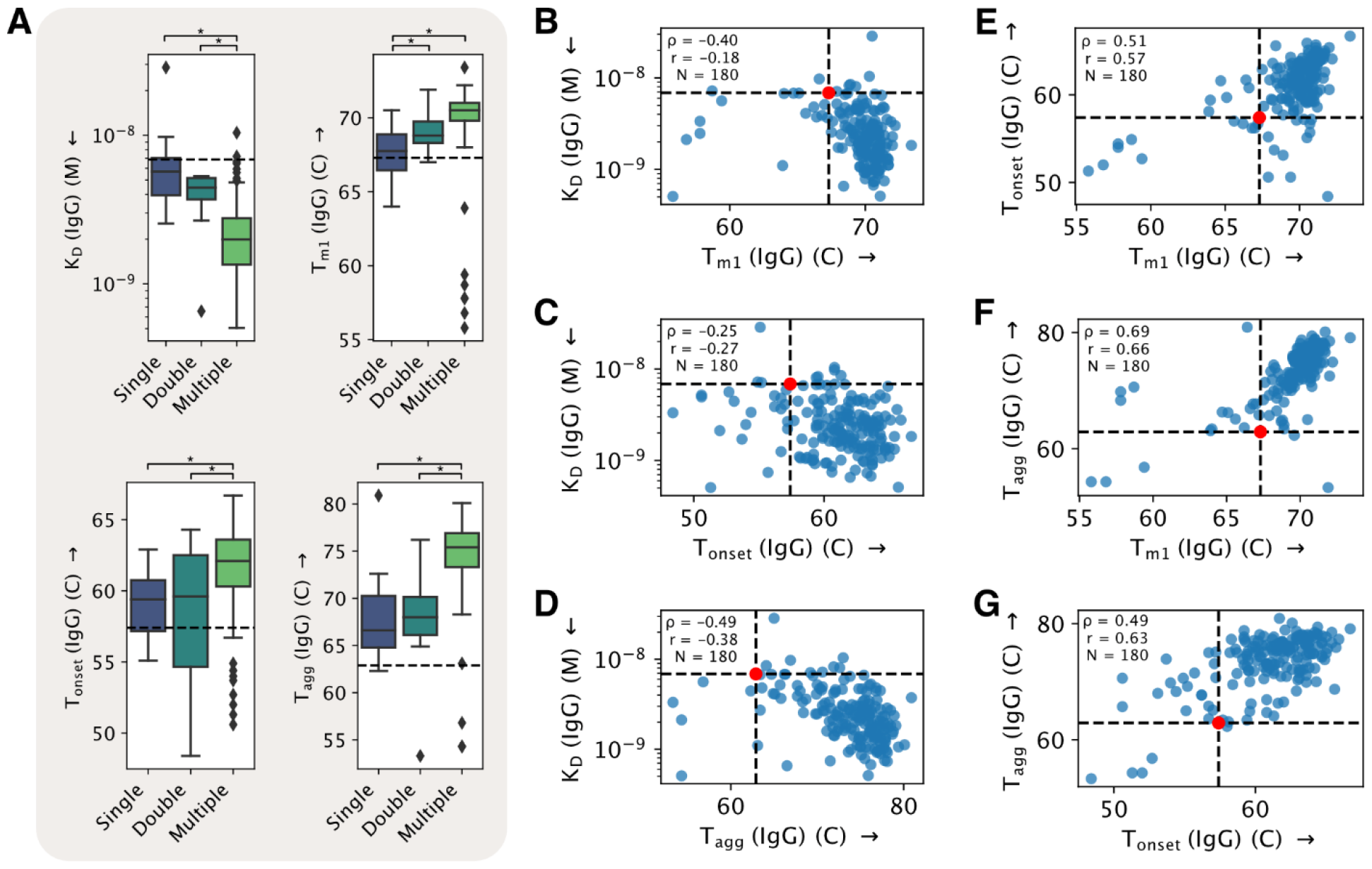
Experimental characterization of “Top 200” optimized variants in IgG format. The D44.1 parental antibody and 199 of the designed variants with the highest DeepAb scores were characterized for thermostability and apparent affinity. (A) Comparison of binding affinity (IgG K_D_) and thermostability (T_m1_, T_onset_, and T_agg_) for optimized variants according to mutational load. Values for the parental antibody are indicated by dashed lines. Significant differences between categories according to a twotailed t-test (p<0.05) are indicated above plots by an asterisk. (B-G) In all plots, variants are represented by blue points and parental antibody is indicated by dashed lines and red points. Arrows on axis labels indicate direction of improved fitness. Spearman (ρ) and Pearson (r) correlation coefficients are calculated for all sequences and reported in each plot, along with the number of total points (N). (B-D) Relationships between thermostability measurements and affinity (IgG K_D_) for optimized variants. (E-G) Relationships between different thermostability measurements for optimized variants. Data for individual clones can be found in Table S1.

Generally, the suggested mutations resulted in improvements in thermostability (T_m1,_ 91%) and affinity (IgG K_D_, 94% of variant IgGs) (Figure 7, Table 1, Table S1). When analyzing how the mutational load (single, double or multiple (5 to 7) mutations per variant) affects affinity (IgG K_D_) and thermostability, significant differences were seen between single and multiple mutation load for all parameters, between single and double mutational load for T_m1_ and between double and multiple mutational load for IgG K_D_, T_onset_ and T_agg_ (Figure 7-A). There was no significant correlation between IgG K_D_ and thermostability (T_onset_, T_m1_, T_agg_) (Figure 7 B-D), while weak correlations were seen between the three thermostability parameters (T_onset_, T_m1_, T_agg_) (Figure 7 E-G).

**Table 1.** Categorizing the changes in affinity (IgG K_D_) and thermostability (T_m1_) in the designed D44.1_IgG_ variants. The data shown here summarize the data found in Figure 7 and Table S1.

|  |  |  |
| --- | --- | --- |
|  | Increased<br>Thermostability (T<sub>m1</sub>) | Decreased<br>Thermostability (T<sub>m1</sub>) |
| Increased affinity<br>(IgG K<sub>D</sub>) | 86% | 8% |
| Decreased affinity<br>(IgG K<sub>D</sub>) | 5% | 1% |

### “Top 27” subset of designed variants show improved thermostability and affinity

The initial screen of the “Top 200” variants for affinity (IgG K_D_) was performed by loading all variants in IgG format onto AHC probes and associating with a single concentration of lysozyme. From within the “Top 200” variants we selected a panel of 30 clones (parental D44.1 and 29 designed variants) that were then re-produced at medium scale (>20 mg) in the IgG format and digested into Fab fragments for further analysis. The selected subset of 29 variants included clones with single, double, or multiple mutations that exhibited a range of high apparent affinities in the IgG binding screen and increased thermostability parameters and were deemed of interest for further analysis (Table S2 and Figure S3). Two of the selected variants, M100 and M114, were discarded after high aggregate levels were discovered. The remaining 27 variants were designated “Top 27” and characterized for affinity (Fab K_D_) towards HEL (Figure 8 and Table S2).

**Figure 8.**
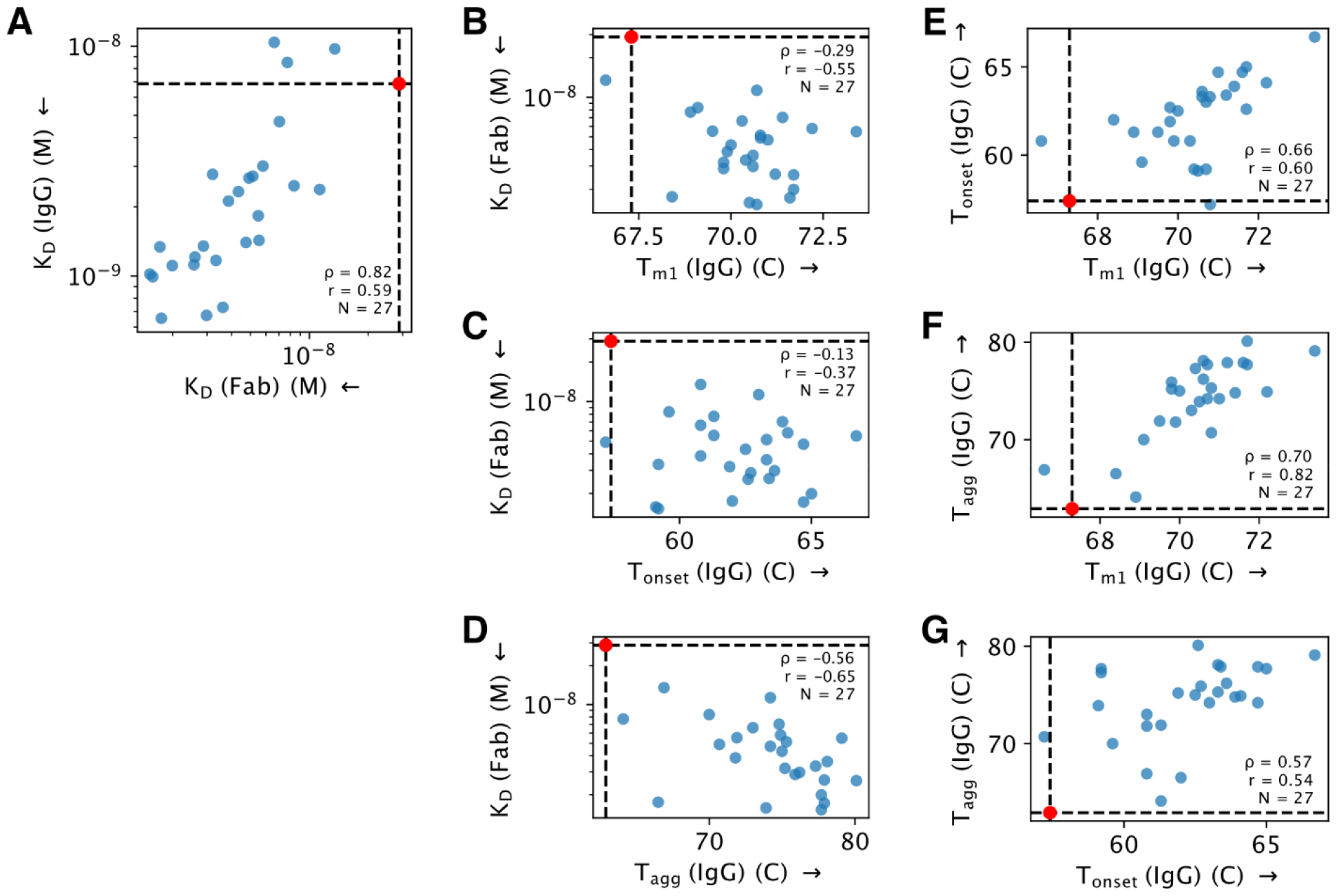
Experimental characterization of the “Top 27” subset of optimized variants. (A) Comparison of binding affinity (K_D_) in IgG and Fab format. Parental antibody binding affinities are indicated by dashed lines. (B-G) In all plots, variants are represented by blue points and parental antibody is indicated by dashed lines and red points. Spearman (ρ) and Pearson (r) correlation coefficients are calculated for all sequences and reported in each plot, along with the number of total points (N). Arrows on axis labels indicate direction of improved fitness. (B-D) Relationships between thermostability measurements and Fab affinity (Fab K_D_) for optimized variants. (E-G) Relationships between different thermostability measurements for optimized variants. Data for the select subset of variants can be found in Table S2.

Fab affinity (Fab K_D_) measurements were performed by loading biotin-conjugated HEL onto SA sensors and associating with dilutions of anti-HEL Fab fragment (the same format that showed weaker Fab affinity previously, Figure 6). This allowed us to alleviate any potential avidity effects on measuring single arm in-solution affinities. All the tested Fabs exhibited improved affinity (Fab K_D_) compared to parental D44.1 Fab (<28.7 nM) (Figure 8-A and Table S2). Improvements in affinity (Fab K_D_) were observed even for single mutations at VL S68 (variants S04 and S10 with affinities of 13.6 nM and 7.7 nM respectively). The largest improvements in affinity (Fab K_D_) were observed for double (D03) and multiple mutations (M017, M018, M022, M117) that exhibited affinities just below 2 nM. These results suggest that mutations at positions VL V57 and S68 are sufficient to improve the affinity to below 2 nM, as mutations at these positions were found in D03 and in all the other highest affinity mutants. IgG affinity (IgG K_D_) and Fab affinity (Fab K_D_) showed weak correlation (Figure 8A).

Re-analyzing the IgG thermostability data variants found in Table S1 only for the “Top 27” subset, most of the variants exhibited increased thermostability (T_m1_, T_onset_ and T_agg_) (Figure 8 and Table S2). Large increases in thermostability (>3 °C increase) were observed for multiple variants: 62% (17/27) of variants showed T_m1_ >70.3 °C, 77% (21/27) exhibited T_onset_ >60.4 °C, and 92% (25/27) exhibited T_agg_>65.9 °C (whereas parental D44.1 exhibits T_m1_=67.3 °C; T_onset_=57.4 °C; T_agg_=62.9 °C) (Figure 8 and Table S2). While improvements in Fab affinity (Fab K_D_) and thermostability do not correlate strongly (Figure 8 B-D), T_agg_ shows a correlation with T_m1_ and T_onset_ shows a mild correlation with T_m1_ (Figure 8 E-G). Overall, 88% (24/27) of the “Top 27” subset showed improvement in both affinity (Fab K_D_) and all three thermostability parameters (T_m1_, T_onset_ and T_agg_).

### “Top 27” subset of designed variants retain the developability profile of parental D44.1

In addition to thermostability and affinity measurements, the developability profile of the “Top 27” subset was assessed in IgG format to ensure that their overall biophysical properties were not adversely affected by the designed mutations (Table 2). The mAbs were tested in a baculovirus particle (BVP) ELISA and a HEK cell binding assay to assess non-specific binding (NSB), AC-SINS to assess reversible self-association (RSA) and accelerated stability heat stress assay to assess aggregation and fragmentation propensity. The expression titers, monomer content after protein A purification, and HP-SEC retention time (RT) were also evaluated.

**Table 2.** Developability profiling of the “Top 27” subset panel of variants. The developability profile of the “Top 27” subset of variants and parental D44.1 in IgG format were characterized using a panel of developability assays. Expression in CHO cells was analyzed by small scale transient transfection compared to the well expressing control mAb NIP228. Monomer content and RT were determined by HP-SEC and compared with the control mAb NIP228. Non-specific binding (NSB) was assayed using a BVP ELISA and a HEK cell binding assay. Reversible self-association (RSA) was assayed by AC-SINS. Aggregation and fragmentation propensity was assessed with an accelerated stability assay conducted at 45°C for 14 days. NT= Not Tested. NA= Not Appllicable

| # | Expression<br>(% of control Ab) | Monomer<br>Content After<br>Protein A<br>Purification (%) | ΔRT from NIP228<br>(min) | BVP ELISA |  | HEK293<br>binding | AC-SINS<br>red shift<br>(nm) | Accelerated stability |  |  |
| --- | --- | --- | --- | --- | --- | --- | --- | --- | --- | --- |
|  |  |  |  | BVP<br>score | BSA/Plate<br>binding<br>(OD 450<br>nm) | Cell<br>binding |  | Change in<br>Monomer<br>(%) | Change in<br>Aggregate<br>(%) | Change in<br>Fragment (%) |
| Parental | 10% | 97.7 | 0 | 1.11 | 0.02 | N | 0 | -1.1 | -0.5 | 1.6 |
| S04 | 54% | 93.9 | 0 | 1.22 | 0.02 | N | 9 | -1.0 | -0.2 | 1.2 |
| S10 | 6% | 98.9 | 0.03 | 1.25 | 0.04 | N | 5 | -4.8 | 4.0 | 0.9 |
| D03 | 101% | 97.4 | 0.11 | 1.05 | 0.02 | N | 10 | -2.2 | 1.5 | 0.7 |
| D05 | 75% | 99.1 | 0.08 | 1.08 | 0.03 | N | 9 | -1.9 | 0.9 | 1.0 |
| M001 | 78% | 97.3 | 0.09 | 1.03 | 0.02 | N | 9 | -1.8 | 0.9 | 0.8 |
| M002 | 72% | 87.9 | 0.1 | 1.65 | 0.02 | N | 0 | -2.5 | 1.7 | 0.8 |
| M013 | 105% | 97.5 | 0.09 | 1.05 | 0.02 | N | 8 | -1.9 | 1.0 | 0.9 |
| M017 | 30% | 96.6 | 0.09 | 2.32 | 0.06 | N | 2 | -2.4 | 1.0 | 1.4 |
| M018 | 94% | 95.8 | 0.1 | 1.37 | 0.02 | N | 9 | -1.6 | 0.7 | 0.9 |
| M022 | 87% | 97.2 | 0.11 | 1.01 | 0.02 | N | 10 | -3.0 | 1.8 | 1.1 |
| M030 | 83% | 95.8 | 0.1 | 1.09 | 0.02 | N | 9 | -1.3 | 0.4 | 0.9 |
| M035 | 12% | 98.8 | 0.08 | 1.3 | 0.03 | N | 1 | -2.9 | 1.2 | 1.6 |
| M039 | 51% | 97.1 | 0.08 | 1.2 | 0.02 | N | 11 | -1.1 | 0.3 | 0.8 |
| M040 | 52% | 96 | 0.07 | 1.04 | 0.02 | N | 0 | -1.0 | 0.3 | 0.8 |
| M041 | 89% | 95.7 | 0.08 | 1.11 | 0.02 | N | 4 | -1.2 | 0.3 | 0.9 |
| M042 | 70% | 98.5 | 0.1 | 0.97 | 0.02 | N | 9 | -1.9 | 0.7 | 1.1 |
| M046 | 73% | 97.2 | 0.08 | 1.08 | 0.02 | N | 1 | -1.7 | 0.9 | 0.8 |
| M048 | 34% | 98.8 | 0.08 | 1.78 | 0.04 | N | 3 | -2.1 | 1.1 | 0.9 |
| M078 | 64% | 97.2 | 0.07 | 4.11 | 0.07 | N | 2 | -1.9 | 1.2 | 0.7 |
| M081 | 11% | 99 | 0.08 | 1.81 | 0.07 | N | 0 | -1.7 | 0.8 | 0.9 |
| M098 | 87% | 97.4 | 0.07 | 1.04 | 0.03 | N | 0 | -2.5 | 1.5 | 0.9 |
| M100 | 12% | 35.4 | NT | NT | NT | NT | NT | NT | NT | NT |
| M102 | 72% | 98.6 | 0.07 | 2 | 0.08 | N | 6 | -2.0 | 0.9 | 1.0 |
| M114 | 19% | 66.9 | NT | NT | NT | NT | NT | NT | NT | NT |
| M117 | 60% | 92.8 | 0.1 | 1.14 | 0.02 | N | 2 | -0.9 | -0.4 | 1.3 |
| M118 | 97% | 96.9 | 0.09 | 1.25 | 0.04 | N | 1 | -2.4 | 1.4 | 1.1 |
| M124 | 40% | 98.1 | 0.1 | 1.41 | 0.04 | N | 5 | -1.8 | 1.0 | 0.8 |
| M157 | 123% | 98.1 | 0.06 | 2.43 | 0.06 | N | 0 | -3.9 | 3.3 | 0.6 |
| M158 | 102% | 98.8 | 0.08 | 1.88 | 0.04 | N | 3 | -2.0 | 1.1 | 0.9 |
| Assay specific positive control | 100% | NA | NA | 68.42 | 1.596 | Y | 36 | NA | NA | NA |
| Assay specific negative control | NA | NA | NA | 1.03 | 0.019 | N | 0 | NA | NA | NA |

Except for one variant (S10), all variants showed increased expression titers compared to parental D44.1, with 72% (21/29) of the designed clones showing a greater than 5-fold increase in expression. Only two variants (M100 and M114) showed extremely low monomer content after protein A purification (35.4% and 66.9%, respectively) and were not tested further. The remaining 86% (25/29) of variants exhibited very high monomer content after protein A purification (>95%) (Table 2). Additionally, all variants had an HP-SEC RT similar to a control IgG1. In the BVP ELISA and HEK cell binding assay, none of the variants showed any significant NSB. In the AC-SINS assay, most variants showed a red shift slightly higher than parental D44.1, but all were much lower than the red shift of 36 nm exhibited by the positive control. In the accelerated stability heat stress assay all the clones showed low levels of fragmentation (similar to parental), while clones S10 and M157 showed an increase in aggregation compared to parental D44.1 (Table 2).

Taken all together, these results show that most of the selected variants exhibit a similar developability profile as parental D44.1. The S68E mutation (variant S10) is clearly deleterious to the yield and colloidal stability, while the exact combination of mutations responsible for the low monomer content of variants M100 and M114 after protein A purification and for the aggregation under heat stress seen for variant M157 are less clear, as these mutations appear in different contexts in other, better-behaved variants.

### Single point mutation variants pinpoint positions responsible for improved characteristics

The mutations found in the designed variants were recreated as single point mutation variants to elucidate the contribution of each mutation in isolation (Table S3). The single point variants where characterized in an identical manner to the previous variants (IgG molecules tested for IgG K_D_ and thermostability then fragmented and purified as Fabs and retested for Fab K_D_). In the point mutants, 42% (9/21) of mutations tested caused an increase in at least one of the thermostability parameters (T_m1,_, T_onset_ or T_agg_), but only three mutations improved two of the parameters (S68M, S68A and N114W) and none improved all three parameters. Additionally, 33% (7/21) of the single point mutations tested caused a >3-fold increase in Fab affinity (Fab K_D_), with 85% (6/7) of the mutations occurring at positions VH N114 or VL V57, and one occurring at position VL S68 (S68E). Improving the affinity (Fab K_D_) to <2 nM seems to require at least two mutations (see variant D03 containing mutations V57A and S68Q, table S2), while achieving improvement in all three thermostability parameters requires 5 mutations (see variants M030 and M040, Table S2).

### Characterization of variants with poor DeepAb scores

Given the significant improvements to thermal stability and affinity observed for the best-scoring variants identified by DeepAb, we next assessed whether low/poorly DeepAb scoring variants would show degradations in these attributes relative to a similar top scoring subset. From the same set of original 4,602 designed variants, we selected 15 low/poor-scoring sequences, each with 5-6 mutations, for characterization and designated them “Low 15 Multi” (Figure 9, Table S4, Figure S4). For comparison we selected the 23 variants with a similar mutational load (5-6 mutations) from our “Top 27” subset and designated these as “Top 23 Multi” (Figure S5). Because even the low-scoring variants (Low 15 multi) were assembled from beneficial DMS mutations, we still expected some improvements over the parental sequence. Indeed, we found that nearly all of the “Low 15 Multi” variants had increased T_onset_ (9/15) and T_agg_ (14/15) compared to parental IgG, but the T_m1_ for most of these variants (10/15) was lower than parental (Figure 9-A and Table S4). All of the “Low 15 Multi” variants exhibited improved Fab affinity (Fab K_D_) compared to parental D44.1 and similar to the comparison “Top 23” subset (Figure 9-A and Table S4). Comparing these two subsets there was no significant difference in Fab affinity (Fab K_D_), but significant increases were seen for T_m1_, T_onset_ and T_agg_ (Figure 9-A). When the subsets were combined to look for correlations, there was no observed correlation between IgG affinity (IgG K_D_) and Fab affinity (Fab K_D_) (Figure 9-B) nor between Fab affinity (Fab K_D_) and thermostability parameters (T_m1,_, T_onset_ or T_agg_) (Figure 9 C-E). Thermostability parameters T_m1_, T_onset_ and T_agg_ did show a correlation with each other (Figure 9 F-H).

**Figure 9.**
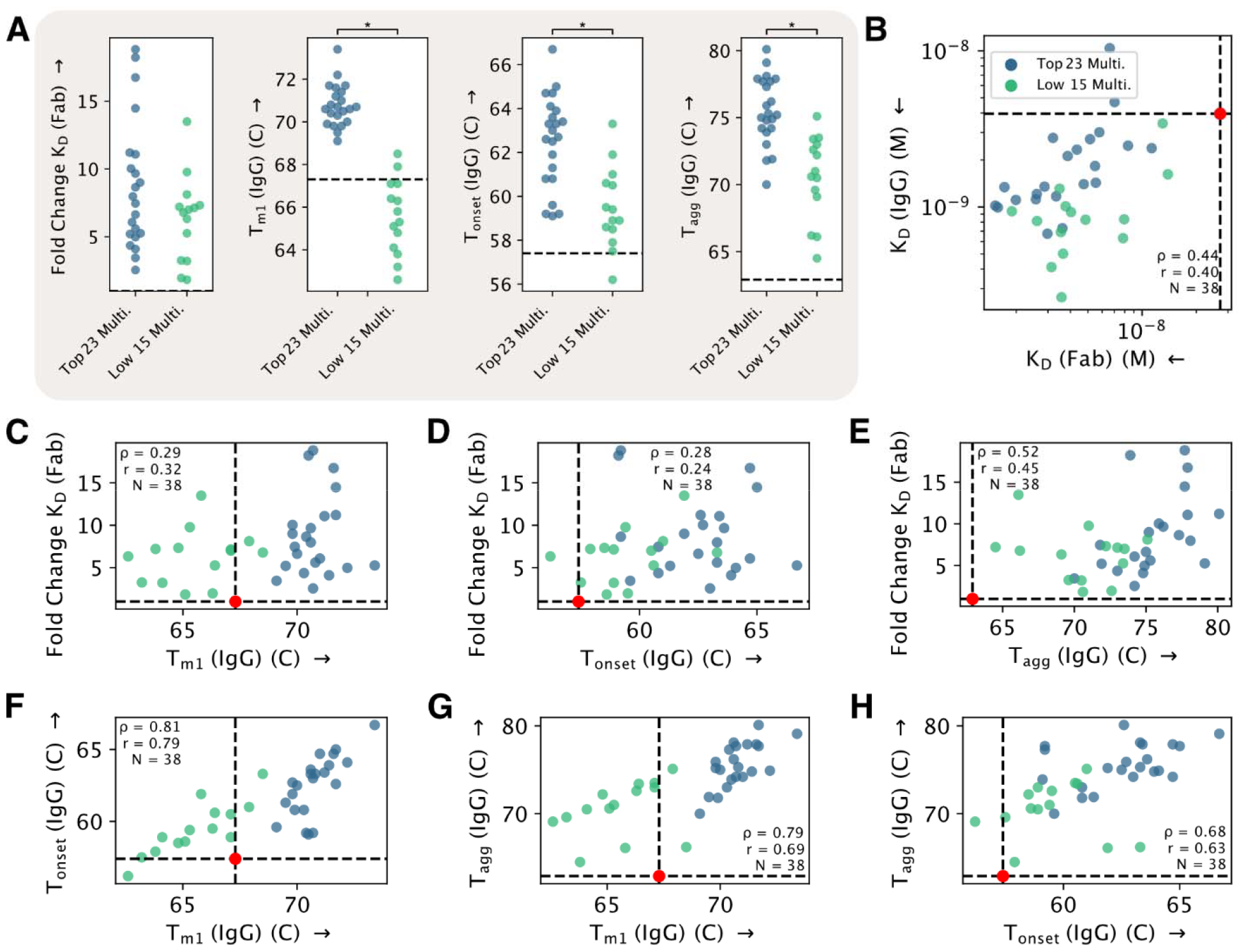
Experimental characterization of “Low 15 Multi” subset of optimized antibodies. The “Low 15 Multi” subset (green), comprised of the 15 lowest/poorest DeepAb scoring variants containing multiple mutations, was characterized and compared to the “Top 23 Multi” subset of optimized variants (blue), comprised of only the variants containing multiple mutations from within the “Top 27” subset of high scoring optimized variants. (A) Comparison of binding affinity (K_D_) and thermostability (T_m1_, T_onset_, and T_agg_) for high scoring and low/poor scoring optimized variants. Values for parental antibody indicated by dashed lines. Significant differences between categories according to a two-tailed t-test (p<0.05) are indicated above plots by an asterisk. (B) Correlation between Fab affinity (Fab K_D_) and IgG Avidity (IgG K_D_). (C-H) In all plots, variants are represented by blue or green points and parental antibody is indicated by dashed lines and red points. Spearman (ρ) and Pearson (r) correlation coefficients are calculated for all sequences and reported in each plot, along with the number of total points (N). (C-E) Relationships between thermostability measurements and binding affinity for optimized variants. (F-H) Relationships between different thermostability measurements for optimized variants.

## Discussion

Antibody structure prediction has rapidly improved in recent years, enabling fast and accurate prediction of variable domains [10, 11, 38]. These advances have been enabled by increasingly accurate deep learning methods trained on experimentally determined antibody structures. Such methods are typically used in antibody engineering workflows to provide structural context for candidate sequences, which can be used to calculate biophysical properties including solubility, aggregation propensity, and thermostability. Prior work has shown that it is possible to improve antibody binding affinity by stabilizing the antibody structure, without direct consideration of the antigen-binding interface [32]. Such methods typically attempt to optimize antibody stability using an atomistic energy function. In this work, we pursued a similar goal, but instead considered structure prediction confidence as a proxy for antibody stability and identified combinations of mutations that yielded the most confidently predicted structure.

An ideal in silico affinity maturation method should operate without the need for any experimental data other than the antibody sequence. By relying solely on antibody sequences, it becomes possible to enhance the affinity or other characteristics of antibodies in the early stages of discovery, even before obtaining any experimental information about the antibody:antigen interface (ie mutagenesis, crystal structures, or cryo-EM). This allows for the optimization of a vast number of antibodies right from the outset. In addition, precise prediction of the antibody:antigen interface is notoriously difficult without additional experimental data, reinforcing the need for sequence based methods. The process we describe here uses an experimental deep mutational scan (DMS) dataset published in Warszawski et al. [32] to target the antibody paratope when designing mutations. Future workflows could instead use DeepAb point mutation scores to identify sites for combinatorial design instead of experimental data.

The combination of DeepAb with our HT (high-throughput) protein production workflow is both versatile and scalable, making it suitable for various engineering objectives. Over multiple iterations, we achieved improvements in both plasmid generation and protein production. Firstly, rapid cloning can be accomplished within 4 days, boasting a success rate of ~90% for identifying the correct sequence in a single round. While this process can be automated at many steps, it can also be carried out completely manually, and therefore should be able to be easily carried out by any lab. Furthermore, through process changes we increased protein production levels significantly, more than doubling the yield achieved with our initial expression and purification process. Again, while the incorporation of full automation in this process can significantly reduce manual intervention and enhance reproducibility, this entire process can also be carried out completely manually without the aid of any robotics automation. The use of this type of HT workflow will significantly benefit not only the research and development of monoclonal antibodies for therapeutic purposes, but also enable the collection of larger datasets to validate proof-of-concept algorithms.

From the 200 antibody candidates proposed by this method, we experimentally determined that 94% exhibited improvements in IgG affinity (IgG K_D_) and 91% exhibited improvements in thermostability (T_m1_) (with 86% exhibiting both increased apparent affinity and thermostability). After screening all candidates for these primary attributes, we selected a subset of 27 clones to further characterize Fab affinity (Fab K_D_) and a series of developability characteristics, including non-specific binding, reversible self-association, expressibility, and aggregation propensity. All of the clones in this subset that were tested for Fab affinity (Fab K_D_) showed improvements in affinity. Additionally, apart from the clones that showed poor expression titers (S10), high aggregation after protein A purification (M100 and M114), or increased aggregation after heat stress (S10 and M157), all remaining clones exhibited favorable developability characteristics as good or better than the parental mAb. Given this data, we find that only a relatively small number (~100) of candidate designs need to be screened to identify mutations that lead to affinity improvements. Depending on the assay, this stage of screening may even be carried out without the need for protein purification, which further improves throughput. Screening for thermostability and developability criteria requires the use of purified protein, but many of the assays described here require using very small amounts of material (i.e., BVP ELISA, AC-SINS, DSF, HP-SEC).

Although we used DeepAb in the present study, the approach developed here is generalizable to any antibody structure prediction model that can produce confidence metrics. Recent methods such as IgFold [10] and ABodyBuilder2 [38] are much faster than DeepAb and provide validated confidence metrics. A similar workflow using either method should enable consideration of more candidate sequences and provide a more faithful proxy for stability. For IgFold in particular, the provided confidence metrics are end-to-end differentiable with respect to the input sequence, meaning the antibody sequence could be directly optimized for stability without the need to computationally screen large combinatorial libraries.

## Materials and methods

### Computing DeepAb score for variants

To prioritize candidate anti-HEL variants for experimental characterization, we compute a mutational fitness score with the DeepAb antibody structure prediction model. Ruffolo et al. defined the DeepAb score as a change in categorical cross-entropy (ΔCCE), which is computed for a given variant relative to the parental sequence as [11]:

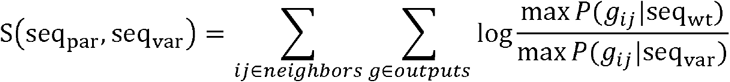

where seq_par_ and seq_var_ are the parental and variant sequences, respectively. Mutated positions in the variant sequence are indexed by *i*. We consider residue pairs *ij*, where *j* is any residue with Cα-distance less than 10 Å from a mutated position *i*. The conditional probabilities represent geometric potentials, *g*, predicted by DeepAb for a particular pair of residues given either the parental or variant sequence. The maximum value of these potentials is taken as a measure of the confidence (or sharpness) of the potential for each sequence. Thus, the log ratio of these values gives a value that measures the change in sharpness upon mutation (lower is better).

### Cloning of anti-HEL Antibodies

Manual high-throughput (HT) cloning was achieved by ordering separate VH and VL DNA sequences in as eBlocks (Integrated DNA Technologies) with proper overhangs for the plasmid of interest and codon optimized for *Homo sapiens*. E-Blocks were cloned using a 4-piece Gibson assembly consisting of the pOE plasmid backbone, VH fragment, VL fragment, and a kappa mid-fragment using Gibson Master Mix (New England Biolabs #E2611L) or HiFi DNA Assembly (New England Biolabs #E2621L) and incubated for 1 hour at 50°C. Gibson assembled plasmids were then transformed into Multishot™ TOP10 Chemically Competent *E.coli* (Invitrogen™ #C4081201) cells using manufacturer’s instructions, in a 96 well format. Transformed cells were then able to be plated in a 96 well format on LB + 100µg/mL carbenicillin and allowed to grow at 37°C overnight. Colonies were handpicked (3 per transformation) and resuspended in LB broth using a 96 well format, these cultures were incubated overnight (800 RPM, 37°C) and miniprepped the following day using a 96 Plasmid NucleoSpin kit (Macherey-Nagel #740625.1). The resulting plasmids were able to be sequenced verified for proper VH and VL insertion. Plasmids were then transferred for HT protein production.

### Cell Culture

Expi293 (Thermo Fisher #A14527) cells were adapted to, grown, and maintained in FreeStyle™ 293 expression medium (Thermo Fisher #12338018) + 1% penicillin/streptomycin (P/S; Thermo Fisher #15140122). Cells were cultured in Erlenmeyer flasks and incubated at 150 rpm in a humidified orbital shaking incubator at 37°C, 8% CO2, 80% humidity. Cells were counted using a ViCell (Beckman Coulter).

### Small Scale Expression in CHO

CHO-G22 cells were maintained in AZ M-030-V8 media (contains 25µM MSX, 100µg/mL hygromycin-B) + 1% penicillin/streptomycin (P/S). Seven days prior to transfection cells were maintained without hygromycin and passaged accordingly. One day before transfection, seed cells at 2E6 VCD at an appropriate volume for experiments in AZ M-030-V2 media + 1% P/S. On the day of transfection cell density was adjusted to be 4E6 VCD, and cells transfected using 3mL of cells, (12E6 cells total), in a 24-well U-bottom plate. Transfection of cells was completed using 18 μL PEI and 3 μg DNA in 50 μL each of 150 mM NaCl. The components were combined for 1 min at room temperature before addition to the cells. Cells were then covered with breathable film and shaken at 250 RPM, 5% CO_2_, 37°C. Cells were fed with proprietary feed after 4 hours and transferred to 34°C incubator, 5% CO_2,_ at 250 RPM. Cells were fed every other day until day 10, where cells were filtered and supernatant titers were quantified using ProA sensors on an Octet Red. The above experiment was repeated in duplicate.

### High Throughput Antibody Expression and Purification

Antibodies were transiently expressed in Expi293 cells using 293fectin™ (Thermo Fisher #12347019) grown in FreeStyle™ 293 expression medium + 1% P/S (Thermo Fisher #12338018). One day prior to transfection, cells were split to 0.7E6 viable cells/mL, or at a density to be 1-1.5E6 viable cells/mL the following day prior to transfection. Just prior to transfection, cells were counted and adjusted to 1E6 viable cells/mL, then plated at 3mL cell suspension/well into 44mm 24-deep-well round-bottom culture plates (Agilent Technologies #202061-300) using a Multidrop Combi (Thermo Fisher).

Plasmid DNA concentration was measured using a Stunner (Unchained Labs). For each transfected well, 2.025µg plasmid DNA and 3µL 293fectin™ were first diluted separately with OptiMEM media (Thermo Fisher #31985070) up to 187.5µL volume and incubated at room temperature for 5 minutes. Plasmid DNA was then added to and mixed with 293fectin™ and incubated at room temperature for 20 minutes before addition to culture wells. Cells were cultured using the Duetz-System of clamps (Kuhner) and covered with 24-well metal sandwich cover lids with black silicone (Enzyscreen #CR1224D). On Day 3 after transfection, cells were fed with 1.5mL fresh FreeStyle™ 293 media and proprietary nutrient supplements.

On Day 7 after transfection, cells were pelleted by centrifugation at 2800 x g for 10 minutes and supernatant was transferred into new 24-deep-well plates, after which 60µL was removed for antibody titer using 384-well tilted bottom plates on an Octet RH16 (Sartorious #18-5080). A 25% slurry of MagSepharose PrismA Protein A magnetic beads (Cytiva #17550001) in 0.1% Tween-20/PBS was added at 200µL/well of supernatant, then incubated on a shaker at room temperature for 4 hours. Beads were washed twice with 900µL 0.1% Tween-20/PBS and twice with 900µL PBS prior to protein elution using 250µL Pierce IgG elution buffer (Thermo Fisher #21009). Eluted proteins were neutralized by adding 30µL 1M Tris-HCl pH 8.0. Neutralized protein concentration was measured using a Stunner (Unchained Labs).

### Thermostability Analysis

The purified antibodies were analyzed by the UNcle system (Unchained Labs, CA, USA) for their thermostability using differential scanning fluorimetry (DSF) and static light scattering (SLS). Briefly, using a laser excitation at 266 nm, intrinsic tryptophan fluorescence and SLS were simultaneously recorded during a linear temperature scan between 25 and 95 °C with a scan rate of 1 °C/min, and with no holding time in order to maximize the frequency of detection. 8.8 µL of each sample (0.5-5 mg/mL) was pipetted undiluted into the UNcle UNI (a sample holding unit containing 16 quartz cells) in duplicates. T_m_ and T_agg_ were analyzed and calculated by the UNcle Analysis Software (V.6.0). UNcle Analysis software determined the T_m_ from the barycentric mean (BCM) of the fluorescence intensity curves from 300–450 nm and the T_agg_ from the intensity of light scattered at 266 nm.

### Fab Fragmentation and purification

IgG in PBS buffer at 1 mg/mL in 10 mM EDTA was added to a solution of equal volume of 2x digestion buffer (40 mM Cysteine in 20 mM Sodium Phosphate, 10 mM EDTA, pH 7.0). Immobilized Papain (Thermo Fisher #20341) was equilibrated with the 2x digestion buffer (500 µL of slurry per 10 mg of IgG). The diluted IgG was incubated with the Papain overnight (16-20 hours) at RT with gentle mixing. The immobilized papain was removed by filtration and the digested protein was dialyzed into 1x PBS, pH 7.2. The Fc portion and uncut IgG was removed by flowing through a 1 mL or a 5 mL MabSelect SuRe column (GE Healthcare #11-0034-95) in PBS (Fab flows through the column). Analysis was completed by quantitating via Nanodrop A280 and using the estimated extinction coefficient of 1.4. Further analysis by SDS-PAGE was completed and aggregation state determined by HPLC-SEC. Molar mass was determined by SEC-MALS.

### Binding kinetics and Quantification by biolayer interferometry (BLI)

Biolayer light interferometry (BLI) was performed using an Octet RED96 instrument (ForteBio; Pall Life Sciences). For measurement of IgG binding kinetics, anti-HEL IgGs were first captured onto AHC biosensor tips (loading 120 s). The IgG loaded biosensor tips then were submerged in binding buffer (kinetics buffer (Sartorius #18-1105) in PBS) containing either single (for the IgG screen) or serial dilutions (for full IgG multi-point kinetics) of HEL protein (Roche #10837059001). For measurement of Fab binding kinetics, biotinylated HEL (HEL_biotin_) was instead captured onto streptavidin (SA) biosensor tips. The HEL_biotin_ coated SA biosensor tips were then immersed in wells containing serial dilutions of anti-HEL Fab. Fits were determined using a 1:1 model. Quantification of IgG titers was performed using Protein A Octet biosensor tips and titers quantified utilizing an established protein titer curve. HEL antigen for the kinetics experiments was purchased (Roche #10837059001) and used either unmodified or biotinylated (using biotinylation reagent Thermo Scientific™ EZ-Link™ Sulfo-NHS-Biotin, #PIA39256 according to manufacturer’s instructions).

### Binding kinetics by Surface Plasmon Resonance (SPR)

The kinetic rate constants (k_a_ and k_d_), and equilibrium dissociation constants (K_D_) of a-HEL IgG for lysozyme was determined at 25°C by SPR on a Biacore T200 instrument (Cytiva, Marlborough, MA). Anti-HEL IgG or a-HEL Fab was immobilized on a CM5 sensor chip with a final surface density of 1000 resonance units (RUs). A reference flow cell surface was also prepared on this sensor chip using identical immobilization protocol but without the a-HEL IgG or a-HEL Fab protein. Two-fold serial dilution of the analyte (lysozyme, 0.78nM to 200nM) was prepared in instrument buffer (HBS-EP buffer; 0.01M HEPES, pH 7.4, 0.15M NaCl, 3mM EDTA, and 0.05% P-20). A single concentration of the analyte was injected over both capture and reference surfaces for 250 seconds at a flow rate of 50µL/minute. The resulting binding response curves yielded the association phase data. Following injection of analyte, the flow was then switched back to instrument buffer for 17 minutes to permit the collection of dissociation phase data, followed by a 60-second pulse of 10mM glycine pH 1.5, to regenerate the analyte-captured surface on the chip. Binding responses against the a-HEL IgG or a-HEL Fab surface was recorded from duplicate injections of each concentration of analyte. In addition, several buffer injections were interspersed throughout the injection series. Select buffer injections were used along with the reference cell responses to correct the raw data sets for injection artifacts and/or nonspecific binding interactions, commonly referred to as “double referencing”. Fully corrected binding data were then globally fit to a 1:1 binding model (Biacore T200 Evaluation software version2.0, Cytiva, Marlborough, MA). These analyses determined the *k*_*a*_ and *k*_*d*_, from which the K_D_ was calculated as *k*_*d*_/*k*_*a*_.

### Size exclusion chromatography

MW analysis of the commercial HEL samples was conducted by UPLC-SEC while developability profiling SEC analysis to determine monomer content was performed by HPLC-SEC. For commercial HEL MW comparison, commercial HEL from an additional source was purchased (Gentex GTX82960-pro) and analyzed by UPLC as described below and utilizing a SEC standard (Biorad #1511901).UPLC-SEC: Samples (100 μg in PHS pH 7.4 buffer, sterile and 0.22 um filtered) were injected in an ultraperformance liquid chromatography (UPLC) Agilent 1290 Infinity II LC system and separated using a 150 mm Agilent AdvanceBio SEC 2.7 micron HPLC Bio size-exclusion column (Agilent #PL1580-3301). The mobile phase used was 100 mM Sodium Phosphate, 100 mM Sodium Sulfate, 0.05% w/w Sodium Azide, pH 7.0, dissolved in Fisher Scientific ultra-pure HPLC grade water (Fisher #7732-18-5). Method flow rate was set to 0.45 mL/min HPLC-SEC: Samples (100 μg in PBS buffer) were injected on an Agilent 1200 series highperformance liquid chromatography (HPLC) instrument and separated using a TSKgel G3000SWxl size-exclusion column (Tosoh Bioscience #08541). The mobile phase was 100 mM sodium phosphate, 100 sodium sulfate, pH 6.8, and sample flow rate was 1 mL/min.

## SDS-PAGE

Purified antibodies were analyzed by SDS-PAGE for aggregation and fragmentation/ purity as HP-SEC analysis would require a larger amount of material. 4 µg of protein was added to a mixture of 2 µL of reducing agent (Intivtogen NuPage #NP004), 5 µL of loading dye (Intivtogen NuPage #NP007) to a final volume of 20 µL using water. Samples were boiled for 10 mins and allowed to cool prior to loading into a 4-12% Bis-Tris gel ((Intivtogen NuPage #NP0321) and run at 200 volts for 30 mins in MOPS buffer (Intivtogen NuPage #NP001). Gels were then washed 3x with water and stained using SimplyBlue SafeStain ((Intivtogen #465034) for 1 hour and destained in water overnight. Gels were then imaged on a Bio-Rad Gel Doc EZ-imager and banding size determined referenced to SeeBlue Plus 2 Presained Sandard ((Intivtogen #LC5925).

### Accelerated Stability Heat Stress Study

Test antibodies were diluted to 1 mg/mL in PBS, pH 7.2 and incubated for 2 weeks at either 4°C or 45°C. Samples were then analyzed by HP-SEC as described above. The monomer, aggregate, and fragment percentages for each antibody were calculated based on curve integration using the HPLC ChemStation software (Agilent). The change in monomer, aggregate, and fragment content was calculated from the difference between each antibody incubated at 4°C versus 45°C.

### Control antibodies

NIP228 is a monoclonal antibody against 4-hydroxy-3-iodo-5-nitrophenylacetic acid67 that historically has been shown not to exhibit aggregation/fragmentation propensities under similar heat stress conditions and is used as a negative control in select developability assays. Control mAb 2 is a monoclonal antibody against an undisclosed target that shows high BVP scores and plate binding in the Baculovirus ELISA and high HEK293 cell binding in the HEK binding assay and is used as a positive control antibody in these assays. “RSA +” control mAb is antibody MEDI1912 and “RSA -” control mAb is MEDI1912_STT (both described in [39]), with high and low RSA scores in AC-SINS respectively. Unless otherwise stated all control antibodies are on a WT human IgG1 Fc backbone. All control antibodies were discovered and produced in-house.

### HEK Binding Assay

Nonspecific HEK cell binding was measured using a Mirrorball Fluorescence Cytometer (SPT Labtech). First, 10 μL of Alexa Fluor 647 goat anti-human IgG (H + L) reporter antibody (Invitrogen #A-21445) was diluted to 16 nM in Mirrorball buffer (Hanks’ Balanced Salt solution with 0.5% BSA) then added to wells of a 384-well, clear bottom plate. Next, each test antibody was serial diluted in Mirrorball buffer and 10 μL of each dilution was mixed with the reporter antibody. Finally, 20 μL of HEK293f cells diluted to 250 cells/μL in Mirrorball buffer were added to the wells. The plate was incubated at room temperature for 2 h, and the fluorescence of each well was measured using the Mirrorball Fluorescence Cytometer. The above experiment was repeated in triplicate

### Baculovirus ELISA

A 1% BV suspension in 50 mM sodium carbonate buffer (pH 9.6) was used to coat half of 96-well ELISA plates (Nunc Maxisorp Cat# 423501) overnight at 4°C, while the second half of the ELISA plates was left uncoated to test the antibodies for BSA/plate binding. All following steps were performed at room temperature. The next day the wells were washed with PBS and then incubated with blocking buffer (PBS with 0.5% BSA) for 1 h, followed by three washes with PBS. The test antibodies were diluted in blocking buffer to concentrations of 100 nM and 10 nM then added to both the BVP coated and uncoated wells for a 1 h incubation followed by three washes with PBS. Goat anti-human IgG-horseradish peroxidase (HRP) secondary antibodies (1:5000 dilution, Sigma-Aldrich #A0170) in blocking buffer were applied to each well and incubated for 60 mins, followed by three washes with PBS. Finally, 3,3’,5,5’-tetramethylbenzidine substrate (SureBlue Reserve, KPL #5120-0081) was added to each well and incubated for 3 mins. The reactions were stopped by adding 50 µL of 0.5 M sulfuric acid to each well. The absorbance was read at 450 nm on a 96-well plate reader (Envision). The BVP scores and plate binding were determined by normalizing absorbance to control wells with no test antibody. The above experiment was repeated in triplicate.

## AC-SINS

Briefly, both whole goat IgG (Jackson ImmunoResearch #005-000-003) (non- capture) and polyclonal goat anti-human IgG Fc (Jackson ImmunoResearch #109-005-098) (capture) antibodies were transferred into 20 mM potassium acetate (pH 4.3) buffer, and then conjugated to 20 nm gold nanoparticles (Innova Biosciences #3201-0100) at a 3:2 ratio of capture:non-capture antibodies. Antibodies were incubated with gold nanoparticles at a 9:1 ratio for 1 h at room temperature, and then blocked by the addition of 0.1 μM poly-ethylene glycol methyl ether thiol (2000 MW, Sigma-Aldrich #729140) for 1 h. The coated and blocked nanoparticles were concentrated by centrifugation to an OD_535_ of 0.4 and stored at 4°C. To assess self-association, 5 μL of nanoparticles were mixed with 45 μL of purified antibody at 50 ug/mL in PBS, pH 7.2 or HA, (20 mM Histidine, 200 mM Arginine) pH 6 in a 384-well plate. Nanoparticles were mixed with buffer only (no antibody) as a control. Absorbance was measured on a SPECTROstar Nano UV/ vis plate reader from 490 to 700 nm. The wavelength of peak absorbance was calculated in the MARS data analysis software and used to determine the wavelength shift compared to the nanoparticle-only control. The above experiment was repeated in triplicate.

### Structural visualization

Structural visualization was conducted using Molecular Operating Environment (MOE) software version 2023.12.

### Code availability

The DeepAb code including a script for design calculations based on (score_design.py) is available at https://github.com/RosettaCommons/DeepAb.

