## Supplemental Figures for "Enhancement of antibody thermostability and affinity by computational design in the absence of antigen"

### Supplementary Figures

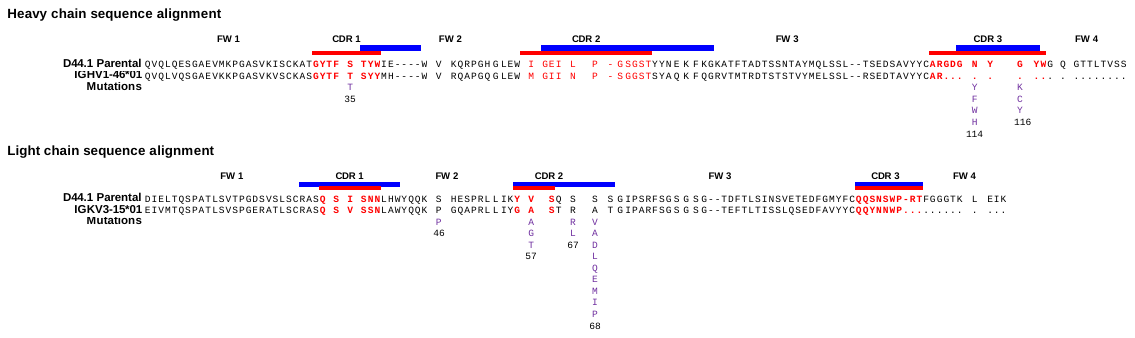

**Figure S1: Sequence analysis of the designed variants.** The sequence of parental anti-HEL D44.1 antibody was aligned against its closest human germlines (IGHV1-46*01 and IGKV3-15*01) and is shown with the mutations used in the DeepAb designed variants marked below the parental sequence. Alignments were conducted using in house BLAZE software. Residues in red font and red bars mark IMGT defined CDRs while the blue bars mark Kabat defined CDRs.

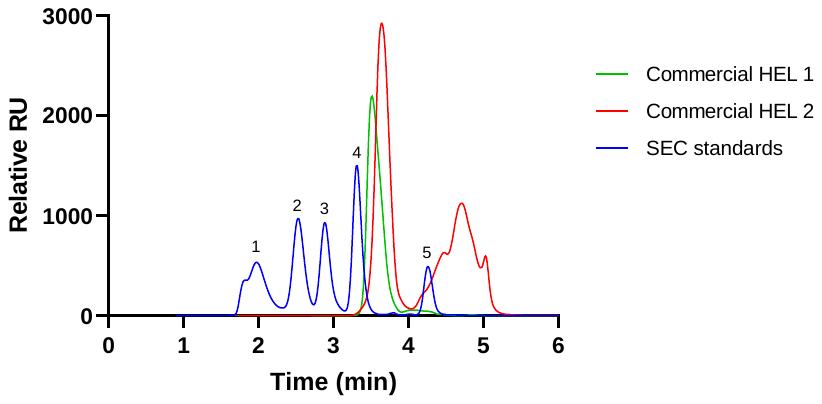

**Figure S2: SEC analysis of commercial HEL.** UPLC-SEC analysis of commercial HEL antigen from either Roche (commercial HEL 1, green trace, used for binding kinetics measurements) or GeneTex (commercial HEL 2, red trace). Both samples exhibit similar main peak elution profiles of the correct size (between standard peaks 4 and 5). The GeneTex sample showed additional peaks (corresponding to sizes below 100 Da when compared to the SEC standards) which are likely buffer components. SEC standard peaks are 1) Thyroglobulin (660kDa); 2) IgG (150kDa); 3) BSA (66.4kDa); 4) Myoglobulin (17kDa); 5) Uracil (0.1kDa).

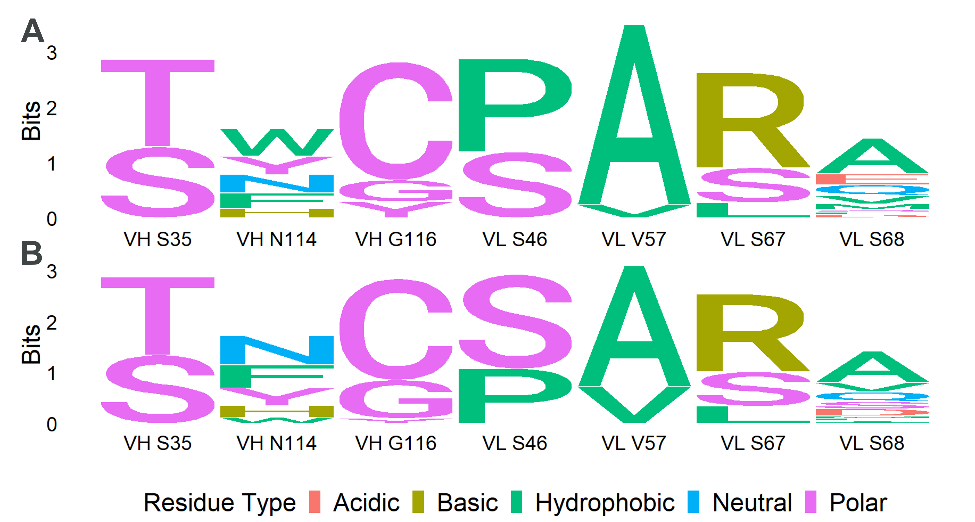

**Figure S3. Sequence Logos for select subset of D44.1. variants.** (A) Subset of 30 antibodies with greatest apparent affinity. (B) Subset of 30 antibodies with greatest thermostability. These subsets are separate, yet overlapping, with the “Top 27” subset that was selected based on thermostability and apparent affinity. Figure was generated using R package ggseqlogo (R version 4.1).

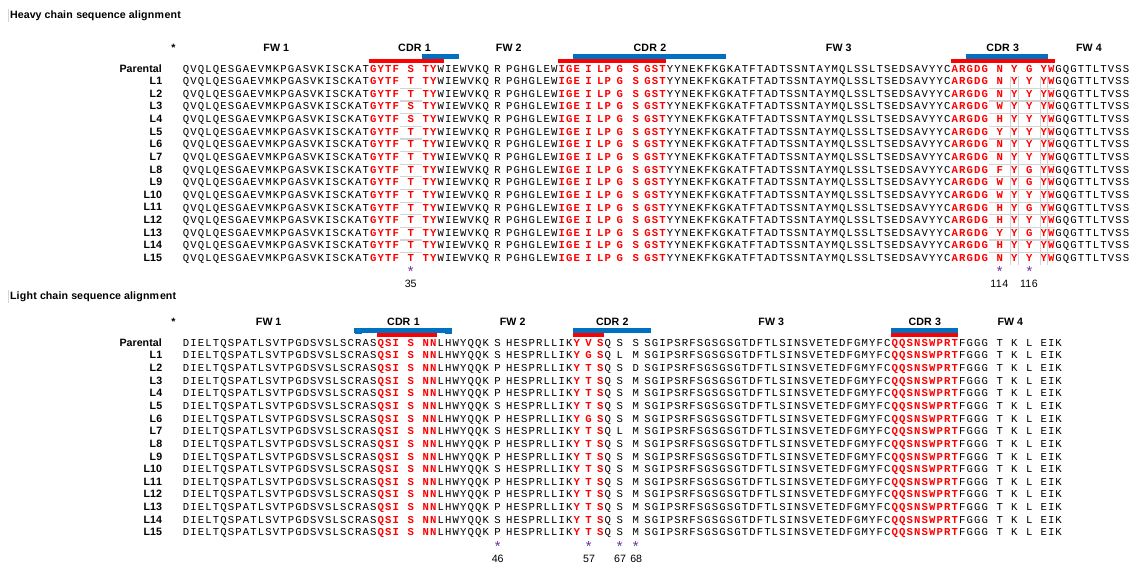
 **Figure S4. Sequence alignment of the parental anti-HEL D44.1 antibody against the 15 antibody sequences with the poorest DeepAb score.** Alignments were conducted using in house BLAZE software. Residues in blue font and blue bars mark Kabat defined CDRs while the red bars mark IMGT defined CDRs. Numbering for the mutations follows IMGT conventions.

-
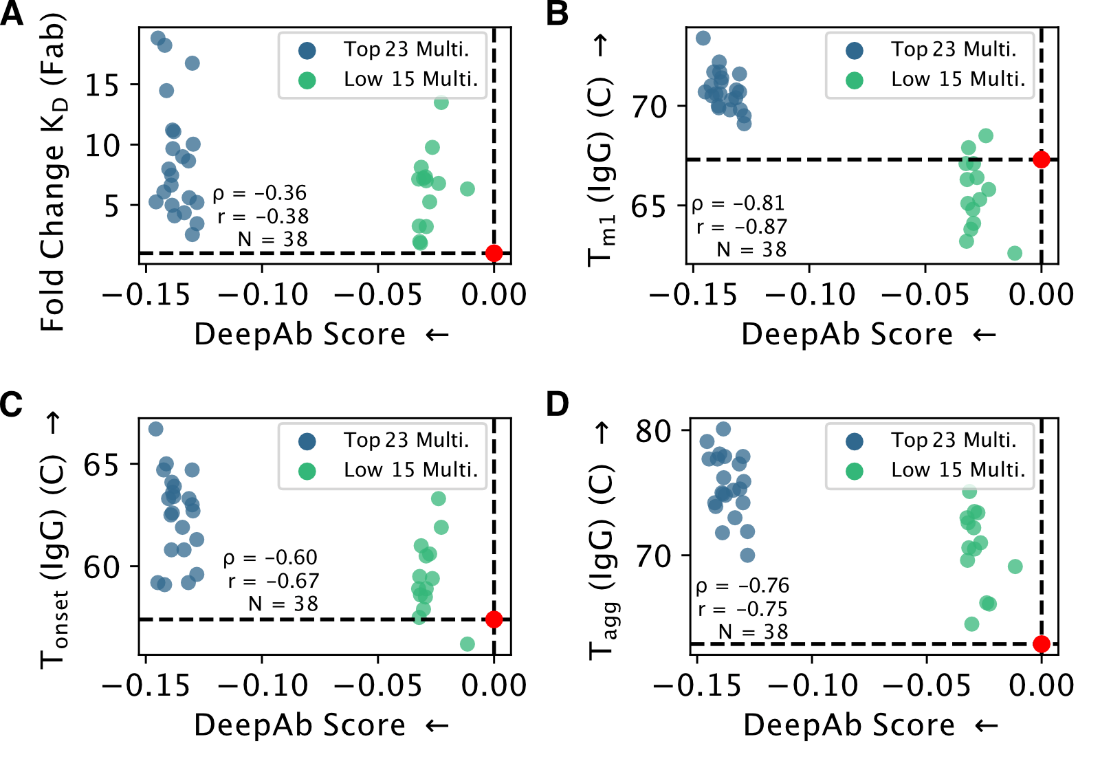

**Figure S5. Relationship between DeepAb score and variant binding affinity (Fab) and thermostability parameters (IgG).** In all plots, variants are represented by blue and green points and parental antibody is indicated by dashed lines and red point. Spearman (ρ) and Pearson (r) correlation coefficients are calculated for all sequences and reported in each plot, along with the number of total points (N). (B) Correlation between Fab Affinity and IgG Avidity. (A) Relationships between DeepAb score and binding affinity for optimized variants. (B-D) Relationships between DeepAb score and different thermostability measurements for optimized variants.
